# Rapid molecular species identification of mammalian scat samples using nanopore adaptive sampling

**DOI:** 10.1101/2023.06.12.544605

**Authors:** Lexi E. Frank, Laramie L. Lindsey, Evan J. Kipp, Christopher Faulk, Suzanne Stone, Tanya M. Roerick, Seth A. Moore, Tiffany M. Wolf, Peter A. Larsen

## Abstract

Accurate species identification is essential to mammalogy. Despite this necessity, rapid and accurate identification of cryptic, understudied, and elusive mammals remains challenging. Traditional barcoding of mitochondrial genes is standard for molecular identification but requires time-consuming wet-lab methodologies. Recent bioinformatic advancements for nanopore sequencing data offer exciting opportunities for non-invasive and field-based identification of mammals. Nanopore adaptive sampling (NAS), a PCR-free method, selectively sequences regions of DNA according to user-specified reference databases. Here, we utilized NAS to enrich mammalian mitochondrial genome sequencing to identify species. Fecal DNA extractions were sequenced from nine mammals, several collected in collaboration with Minnesota Tribal Nations, to demonstrate utility for NAS-barcoding of non-invasive samples. By mapping to the entire National Center for Biotechnology Information (NCBI) mammalian mitochondrial reference genome database and bioinformatically analyzing highly similar matches, we successfully produced species identifications for all of our fecal samples. Eight of nine species identifications matched previous PCR or animal/fecal morphological identifications. For the ninth species, our genetic data indicate a misidentification stemming from the original study. Our approach has a range of applications, particularly field-based wildlife research, conservation, disease surveillance, and monitoring of wildlife trade. Of importance to Minnesota tribes is invasive species monitoring, detections, and confirmation as climate impacts causes changes in biodiversity and shifts in species distributions. The rapid assessment techniques described here will be useful as new introductions and range expansions of native and invasive species may first be detected by the presence of signs such as scat rather than direct observations and will be helpful for chronically understaffed tribal natural resources agencies.

## Introduction

Species identification is essential to the study of mammals, especially for ecological, biodiversity, and conservation-based studies. Nevertheless, accurate identification can be difficult, especially between cryptic, elusive, and endangered species that are rare to observe in nature or within natural history collections. An array of methods, with varying levels of accuracy, are routinely used to identify mammalian species ranging from direct field observation and examination of external morphological characteristics to morphometric analyses of subtle cranial features and molecular barcoding using DNA. Of these methods, molecular approaches that facilitate the phylogenetic, phylogeographic, and molecular systematic analyses of mammals have led to a large increase of the recognized mammalian species in nature, especially among taxa of bats, rodents, and shrews (Bradley and Baker 2001; Redondo et al. 2008; Baird et al. 2015; Giarla and Esselstyn 2015; D’Elía et al. 2019; DeSalle and Goldstein 2019).

Advancements in molecular technologies and associated methodologies continually provide new research opportunities for the study and identification of mammals. For example, the usage of fecal samples for host identification. Feces is an easy to acquire, non-invasive sample that can be used to elucidate many biological features of the depositing host species (genetics, diet, metacommunities) without needing to handle or visualize the animal directly (Kohn and Wayne 1997; Valentini et al. 2009; Srivathsan et al. 2016; Kusack et al. 2022; Pannoni et al. 2022). Although morphological examination of mammalian feces, including size and shape, are routinely used to distinguish species, even subject matter experts can fail to produce an accurate species identification from fecal morphology alone (Davison et al. 2002). More recently, molecular techniques using fecal DNA have proven useful to achieve more accurate identifications. Sloughed rectal cells from the excreting individual are present in the feces and can be used to extract host-specific DNA (Höss et al. 1992). An early and still widely used approach consists of PCR amplification of host barcoding genes, typically the mitochondrial cytochrome-*b* (cytb) and/or cytochrome oxidase I (COI) gene for mammals, and sequencing of fecal DNA, with a wide variety of downstream applications ranging from phylogenetics to forensics. For example, this method was used to investigate the genetics of a threatened bear population in Europe that consisted of less than 10 individuals spread across a large geographic area (Höss et al. 1992). In another study, species-specific primers were used to identify feces from the elusive and rare *Lynx pardinus (*Iberian lynx), however low concentrations and quality of DNA generated false negatives (Palomares et al. 2002). In a similar study, researchers identified species from morphologically similar fecal samples of multiple sympatric carnivores by utilizing a multiplexed PCR system that produced fragments of different lengths for each species (Dalén et al. 2004). However, this method could only account for species included in the specific set of primers targeted for the study and was challenged by false negatives.

Collectively the utility of PCR and specific primers for the identification of mammal species from scat suffers from several limitations. Despite the ease of collection, fecal samples are composed of a complex mix of biological material, including bacteria, that can inhibit downstream molecular techniques, especially PCR (Kohn et al. 1995). The presence of bacteria, the initial warm and damp environment of feces, and subsequent environmental exposure leads to degradation of host DNA (Kohn et al. 1995). Despite the extensive usage of scat-based PCR and sequencing of host barcoding genes, the method is met with significant challenges, including time, expense, need for species-specific primers, limited quality and quantity of DNA, and false negatives. Moreover, current PCR-based molecular techniques for scat-based identification largely require molecular-grade laboratory conditions, meaning that time to identification can be substantial and the techniques are typically not performed in the field.

Genomic technologies have advanced rapidly over the past few decades and are impacting the field of mammalogy in remarkable ways (Larsen and Matocq 2019). One of the most exciting advancements pertains to single molecule nanopore sequencing, with a variety of sequencing platforms and applications introduced by Oxford Nanopore Technologies (ONT). Since 2014, the per-base accuracy of ONT sequencing platforms has steadily improved, with current raw sequence rates achieving greater than 99% accuracy per base (Oxford Nanopore Technologies 2023). When combined with the field-deployable ONT MinION sequencer, such improvements open the door to a wide variety of applications for field-based molecular research. In parallel to ONT hardware and sequencing chemistry improvements, significant advancements have been made in bioinformatic algorithms for the real-time analyses of ONT sequence data. In particular, the recently released nanopore adaptive sampling (NAS) software (i.e., ReadUntil) can be used to selectively sequence individual molecules of DNA, cDNA, or RNA (Payne et al. 2021; Martin et al. 2022; Kipp et al. 2023).

As individual molecules are being sequenced within a given nanopore, NAS utilizes an advanced bioinformatic pipeline (e.g., minimap2) to compare the resulting nucleotides to a user-specified reference file (e.g., all publicly available mammalian mitochondrial genomes), with real-time results generated during a given sequencing experiment. Approximately every 0.4 seconds, a sequence of a given molecule is compared to the reference. Any sequences having a minimum of ∼70% similarity to the reference database will be retained and those that are below ∼70% similarity to the database are rejected (Payne et al. 2021). Therefore, targets of interest can be selectively enriched (e.g., mitochondrial DNA (mtDNA), specific genes, pathogen genomes, etc.) and non-target DNA is rejected from the sequencing pore. NAS can effectively be applied for host-species identification of fecal samples by rejecting non-mammal DNA and enriching for putative host DNA (Figure 1)(Payne et al. 2021; Wanner et al. 2021). In particular, the method is ideally suited for the targeted enrichment of mammalian mtDNA, present in high copy numbers in cells which helps avoid obstacles associated with degraded samples or highly repetitive DNA (Wanner et al. 2021). An important aspect of NAS is that the length of a sequenced read is not restricted to template length and single-molecule sequences can be thousands of bases long.

**Fig. 1.**
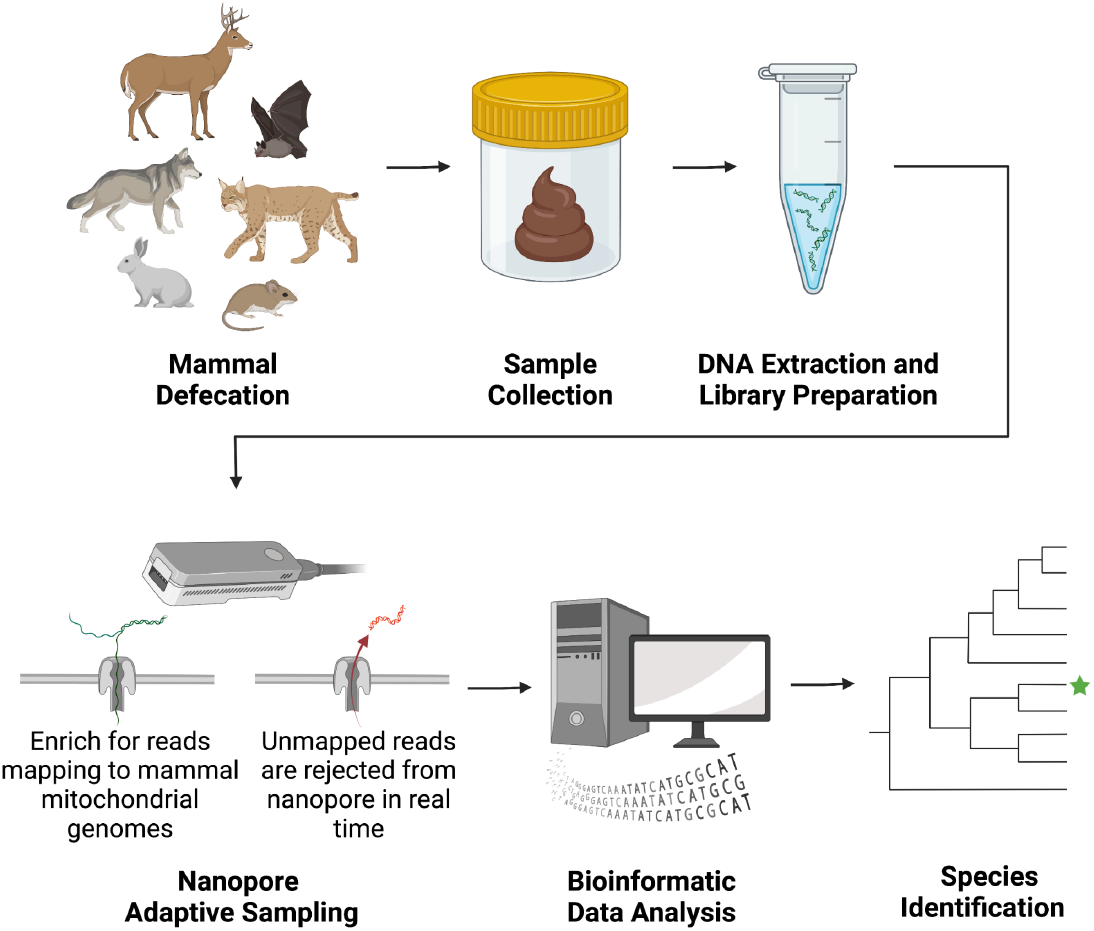
Species identification pipeline using Nanopore Adaptive Sampling. Mammal feces is collected from the environment and whole genomic DNA is extracted. DNA extracts are prepared for Oxford Nanopore Sequencing following protocols for genomic DNA. NAS enriches for sequencing of mammalian mitochondrial genomes. Sequenced reads are bioinformatically analyzed to determine species. Created with BioRender.com.

Long reads with overlapping sequences of DNA can be used to increase confidence of downstream taxonomic identifications and can potentially recover and be used to assemble entire mitochondrial genomes. Importantly, the methodology we describe does not require PCR to amplify specific barcoding genes (e.g., cytochrome-*b* or COI).

Indigenous nations have long been stewards of biodiversity globally and have interest in maintaining strong stewardship principles using both traditional and modern techniques (Fletcher et al. 2021). Practical applications of rapid species identification technologies will assist tribal and other governmental entities in determination of mammalian species assemblages using non-intrusive techniques such as scat collections and opportunistic field collections. Our aim was to develop a tool to enhance applied field assessments of biodiversity and invasive species monitoring. In this study, we merged modern technology developments in genetic assessments with tribally collected data to evaluate the efficacy of rapid species detection capabilities.

Here we show how the NAS method can be leveraged for the rapid sequence-based molecular identification of a variety of mammalian species using DNA extracted from scat samples. Using purely a bioinformatic approach, NAS can be used for real-time mammalian species identification without PCR. Of particular interest to the research community is that the method can be performed with field-deployable equipment using a straightforward whole genomic DNA extraction and sequencing approach with mapping results reported during the analysis, thus facilitating rapid putative species identifications in both lab and field settings.

## Materials and Methods

### Sample Collection and DNA extraction

Fecal samples were collected via a previous study conducted across several sites including from three Minnesota Ojibwe tribes in northern Minnesota, the Leech Lake Reservation, the Red Lake Reservation, and Grand Portage Reservation (Bernstein et al. 2021) as well as for ongoing research efforts. All activities have been reviewed and approved by the University of Minnesota IACUC [protocols 1803-35736A; 1809-36374A] and all methodologies involving live animal handling follow ASM guidelines (Sikes and the Animal Care and Use Committee of the American Society of Mammalogists 2016). For samples originating from Bernstein et al. (2021; see Table 2), species identification was originally confirmed by a multiplex polymerase chain reaction (PCR) that amplified fragments of mtDNA from the control region (De Barba et al. 2014) or the sample was taken directly from a known species. All other samples were opportunistically collected in Minnesota and identified either by fecal morphology to the lowest taxonomy possible or by morphological characteristics of the excreting mammal at the time of collection. These species identifications were blinded for the NAS analysis and revealed for comparison after NAS-guided species identification was completed. Samples were stored at -80°C until DNA extraction was performed. DNA was extracted from nine samples from a variety of mammal species using the QIAmp® PowerFecal® Pro DNA Kit (QIAGEN®, Hilden, Germany). DNA extracts were quantified using a Qubit 4 fluorometer (Invitrogen, Carlsbad, United States). From each sample, the input DNA concentration going into library preparation ranged between 3.74 and 780 ng/μl.

### Nanopore Library preparation

Library preparation was performed blinded to sample species identification. Nine individual DNA libraries were constructed using the ONT Sequencing Ligation Kit SQKLSK109 or SQKLSK110 following the manufacturer’s instructions for either the R9.4.1 Flongle (up to 126 sequencing nanopores, low throughput) or R9.4.1 full-sized MinION flow cell (up to 512 sequencing pores, high throughput) protocols.

Total input DNA into library preparation from each sample ranged between approximately 500 to 1000 ng, for use with the Flongle or full flow cell, respectively. To prepare the ends of the DNA molecules in each sample for adapter attachment, we used NEBNext FFPE DNA Repair and Ultra II End Repair Module reagents (New England Biolabs Inc., Ipswich, United States) and incubated at 20°C for 5 minutes and then at 65°C for 5 minutes. AMPure XP beads (Beckman Coulter, Indianapolis, United States) were used for bead clean-up on a magnetic separation rack. For adapter ligation, Adapter Mix F (Oxford Nanopore Technologies, Oxford, United Kingdom), Ligation buffer (Oxford Nanopore Technologies, Oxford, United Kingdom), and NEBNext Quick T4 DNA Ligase (New England Biolabs Inc., Ipswich, United States) were added to the DNA sample from the previous steps and incubated for 10 minutes. This was followed by another bead cleanup and addition of the ONT Short fragment buffer to purify fragments of all sizes equally. The final library was eluted into an Elution buffer (Oxford Nanopore Technologies, Oxford, United Kingdom) of volume of 7 μL for Flongle sequencing or 15 μL for sequencing on a full flow cell at 37° C and quantified by a Qubit 4 Fluorometer (Invitrogen, Carlsbad, United States). Each library was stored at 4°C before sequencing.

### Sequencing and Basecalling

Each library was individually sequenced on a MinION with either a R9.4.1 Flongle flow cell or R9.4.1 full flow cell. The sequencing was performed on either a Linux desktop computer (Intel C600/X79 series i9-10920X 12 core; Linux 5.4.0-77-generic x86_64; Ubuntu 18.04; Nvidia Quadro RTX 4000 GPU with 8 GB video memory) or a Linux laptop (16x 11th gen Intel Core i7; Ubuntu 18.04; Nvidia GeForce RTX 3080 Ti GPU with 16 GB video memory). Flush Tether and Flush buffer (ONT) were mixed and loaded to prime the flow cell. The library, Sequencing Buffer II, and Loading beads II (ONT) were combined and loaded into the flow cell. Sequencing parameters were set within the MinKNOW GUI (ONT, v4.3.20) with the adaptive sampling option turned on. The adaptive sampling reference file was created by gathering all reference mammal mitochondrial genomes available from National Center for Biotechnology Information (NCBI) and including them in a single reference file in FASTA format. This file includes all the complete reference mitochondrial genome sequences from all mammal species available on NCBI at the time of the file’s creation (14 March 2022). This file is selected during the setup of an adaptive sampling experiment to determine enrichment of reads and alignment in real-time. Fast basecalling was chosen for the real-time alignments to the reference file. Sequencing was initiated and run for 48 hours or until the flongle/flow cell was exhausted of pores. Raw FAST5 files generated during sequencing were basecalled post-hoc with super accuracy by ONT Guppy basecaller (v5.0.11). Raw FASTQ nanopore sequence data from the experiments are deposited in the NCBI Sequencing Read Archive (SRA), under project number [Accession number pending, provided upon publication].

### Bioinformatic Processing

Detailed instructions describing the pipeline used here are available in Supplementary Data SD1. Bioinformatic analysis was completed with access to the Minnesota Supercomputing Institute, which provided computational resources and data storage. For each sequencing run, metadata was generated using the nanoplot (v1.32.1) and fastqc (v0.11.7) software packages. FASTQ files generated for each sample were concatenated and then quality filtered using a score of 7 or higher and a read length between 300 bases and 17 kilobases using NanoFilt (v2.6.0). This FASTQ was then aligned to the reference file containing all mammal mitochondrial genomes using minimap2 (v2.17). Files were indexed and organized using samtools (v1.9). The filtered FASTQ for each sample with reads mapping to the mitochondrial database was then used as input into Kraken2 (v2.1.2) to further filter the data for mammalian mitochondrial reads. Kraken2 maps the reads to a reference database using a *k*-mer-based approach to provide taxonomic classifications of sequences. Here, the Kraken2 reference database was created using the NCBI mitochondrial genome refseq file. Kraken2 output files were used to visualize the data in Pavian software on R studio (v4.2.2). *De novo* assembly of the data was completed with Flye (v2.9.1), when possible, and contigs were used for phylogenetic analyses. Flye assembly contigs of near-complete mitochondrial genomes were annotated with the web-based annotator, Mitos2 (Donath et al. 2019).

### Phylogenetic Analysis

When possible, barcoding genes (cytb and COI) or any substantial and continuous sections of the mitochondrial genome were extracted from sequenced reads by aligning to matching reference mitochondrial genomes using Geneious Prime software (v2022.2.1). For rapid putative results, these barcoding genes or large sections of mitochondrial sequences were input into the NCBI Basic Local Alignment Search Tool (Blast) search engine and/or the Barcode of Life Data System (BOLD) identification search engine for COI genes.

These web-based search engines provide the top matching sequences from their databases with percent similarity calculations. The Blast search provides ‘Distance tree results’ consisting of a neighbor-joining phylogenetic tree of our sample sequence with the matches generated by Blast. The BOLD search engine provides a ‘taxon ID tree’ that generates a neighbor-joining tree based on our sample and their matching nucleotide sequences.

To confirm these rapid results provided by Blast and BOLD search engines, barcoding genes from the same species and closely related species, as well as an outgroup species, were collected from NCBI (accession numbers provided in figures). Phylogenetic trees were constructed using RaxML (v8.2.12) GAMMA model with 1,000 bootstrap iterations.

## Results

We successfully sequenced mitochondrial sequences and/or particular barcoding genes (cytb and/or COI) from all fecal samples using the NAS method, including the near complete mitochondrial genome of two samples. Putative species identifications of blinded samples were initially based on real-time NAS mapping results. Molecular data was generated for each sample through individual sequencing experiments (Table 1). Depending on flow cell type (i.e., Flongle vs. full-size Minion flow cell), the total number of bases sequenced for each sample ranged from 63,666,885 to 4,909,175,834 bases and total number of reads ranged from 157,425 to 10,502,105 reads. Based on our NAS bioinformatic results, eight out of nine of our identifications matched with previous identifications to species-level after unblinding sample identifications (Table 2).

**Table 1.** Nanopore adaptive sampling sequencing experiments for rapid mammalian species barcoding. Samples collected in Minnesota from wildlife or zoo animals.

| Sample ID | Flow cell type | Sequencing pores at experiment start | Total bases sequenced | Total number of reads | Reads after filtering |
| --- | --- | --- | --- | --- | --- |
| 1 | Flongle | 65 | 605,520,941 | 1,311,441 | 967,924 |
| 2 | Flow cell | 357 | 4,555,260,593 | 10,502,105 | 6,701,295 |
| 3 | Flongle | 80 | 175,525,810 | 433,311 | 264,108 |
| 4 | Flow cell | 219 | 69,091,774 | 296,158 | 56,317 |
| 5 | Flongle | 47 | 273,448,450 | 615,773 | 457,829 |
| 6 | Flow cell | 787 | 4,909,175,834 | 9,786,381 | 6,029,765 |
| 7 | Flow cell | 492 | 2,702,000,834 | 5,136,945 | 4,061,666 |
| 8 | Flongle | 49 | 157,230,291 | 330,846 | 258,833 |
| 9 | Flongle | 23 | 63,666,885 | 157,425 | 111,708 |

**Table 2.** Nanopore Adaptive Sampling and mitochondrial mapping results from mammalian fecal samples leads to species identification. All samples were collected in Minnesota from 2020 to 2022. Fecal morphology IDs are based on identifications made during fecal sample collection. Original species ID are made based on depositing animal morphology or PCR amplicon size. NAS species IDs were generated using the pipeline presented herein. Reads mapped to the mitogenome database indicate the number of nanopore sequencing reads with matches to the NCBI Refseq database. Total number of mapped bases indicates the total number of bases in the reads that mapped to the NCBI Refseq database.

| Sample ID | Fecal morphology ID | Original species ID | NAS species ID | Reads mapped | Total bases mapped |
| --- | --- | --- | --- | --- | --- |
| 1 | - | *Canis lupus familiaris*<sup>a</sup> | *Canis lupus* | 98 | 9,644 |
| 2 | *Canis lupus* | *Canis lupus*<sup>b</sup> | *Canis lupus* | 1744 | 14,425 |
| 3 | *Canis lupus* | *Vulpes vulpes*<sup>b</sup> | *Canis lupus* | 41 | 8,116 |
| 4 | *Vulpes vulpes* | *Pekania pennanti*<sup>b</sup> | *Pekania pennanti* | 47 | 2,102 |
| 5 | *Canis lupus* | *Lynx rufus*<sup>b</sup> | *Lynx rufus* | 53 | 2,845 |
| 6 | *Sylvilagus floridanus* | *Sylvilagus floridanus*<sup>a</sup> | *Sylvilagus floridanus* | 38 | 6,173 |
| 7 | Rodent spp. | Rodent spp.<sup>a</sup> | *Peromyscus leucopus* | 173 | 11,024 |
| 8 | *Odocoileus virginianus* | *Odocoileus virginianus*<sup>a</sup> | *Odocoileus virginianus* | 73 | 6,685 |
| 9 | - | *Eidolon helvum*<sup>a</sup> | *Eidolon helvum* | 24 | 1,079 |

<sup>a</sup> indicates animal morphology-based identification, <sup>b</sup> indicates PCR amplicon size identification.

Kraken2, Blast and BOLD matches, and generation of phylogenetic trees produced aligning species identification results (Figure 2). The BOLD phylogenetic tree that was produced for our sample 8 showed paraphyletic grouping with two closely related species of deer, *Odocoileus virginianus* and *Odocoileus hemionus* (See discussion). We further investigated the sample identification that did not match with the previous PCR identification, sample 3. Sample 3 was identified by fecal morphology as *Canis lupus* (gray wolf), then identified by PCR as *Vulpes vulpes* in Bernstein et al. (2021). All our bioinformatic analyses for this sample provided support to identify the sample as *C. lupus*. We found no evidence to support the *V. vulpes* identification. From two sequencing experiments with full-sized flow cells, sample 2 (*C. lupus*) and 7 (*Peromyscus leucopus*), near complete mitochondrial genomes were assembled with 61 and 24X coverage, respectively (Figure 3).

**Fig. 2.**
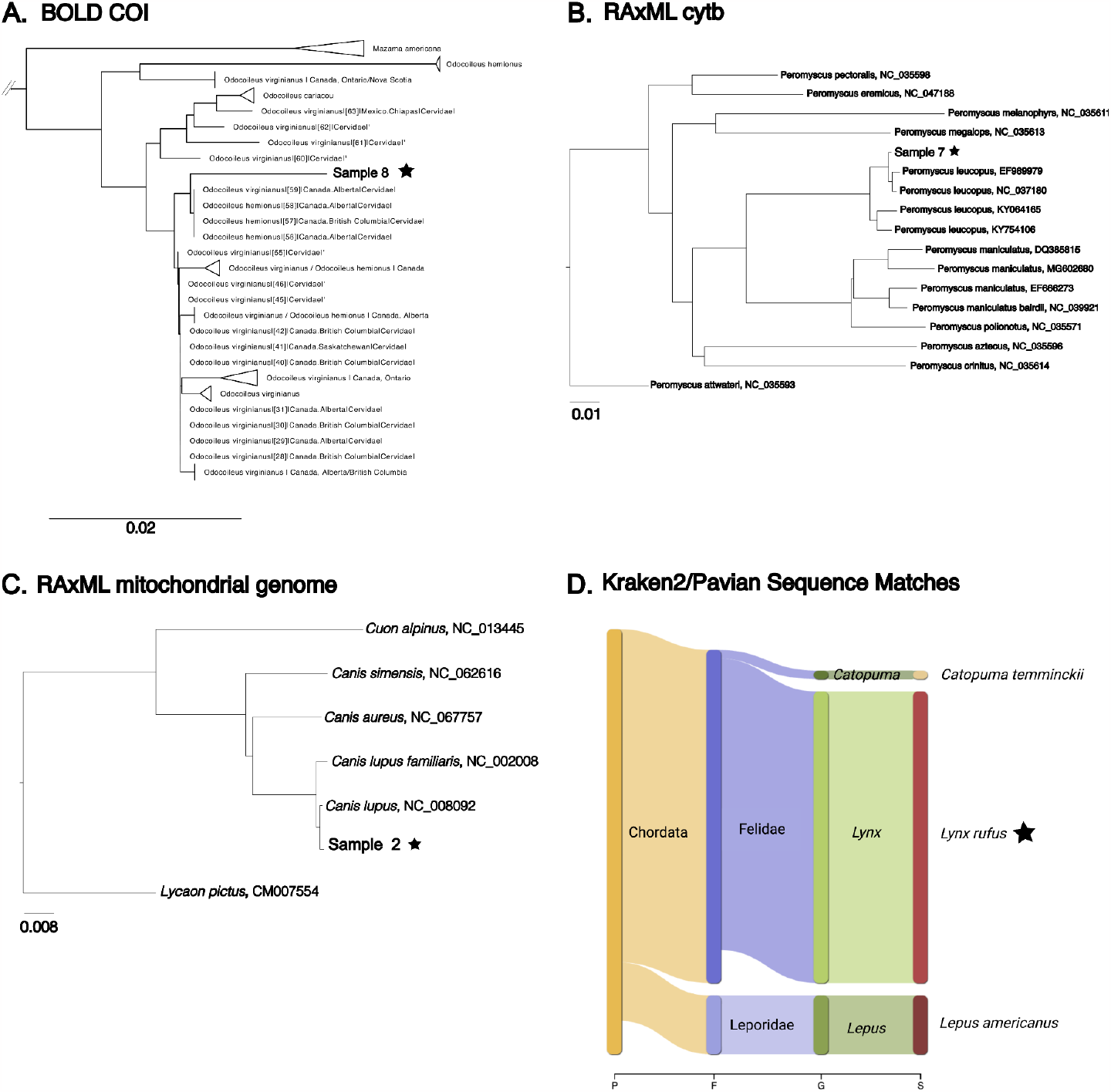
Various bioinformatic methods can be utilized to produce species identification. Our samples or the species identification are indicated with the black star. Panel A shows a neighbor joining phylogenetic tree generated by the BOLD database using COI (accession numbers provided in Supplementary Data SD2). Panel B shows a phylogenetic tree generated by RAxML using cytb sequences from sample 7 and closely related species from NCBI. Panel C shows the full mitochondrial genome of our sample 2 in a RAxML phylogenetic tree with closely related species from NCBI. Panel D shows the Pavian visualization of kraken2 database matches. Reads from sample 5 matched with 3 species, *Catopuma temminckii, Lynx rufus*, and *Lepus americanus*. The majority of the reads matched to *Lynx rufus*.

**Fig. 3.**
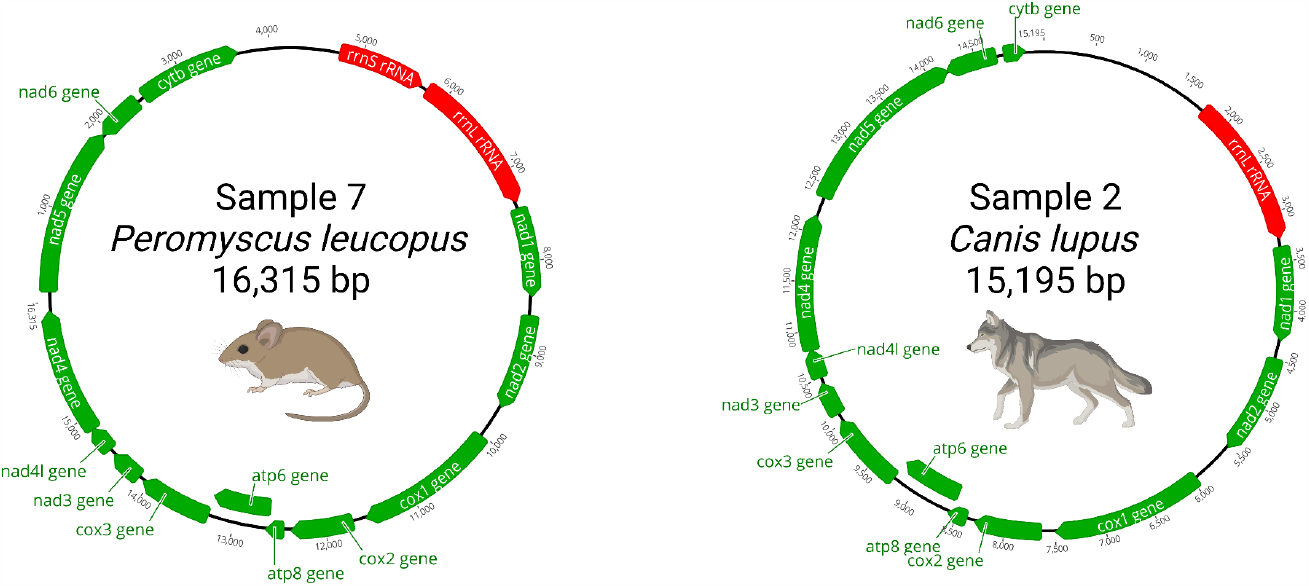
Complete and near-complete mitochondrial genomes from *P. leucopus* (sample 7) and *C. lupus* (sample 2), respectively. Both mitochondrial genomes were sequenced using nanopore adaptive sampling. *C. lupus* mitochondrial genome is near-complete, with approximately 911 bps missing from cytb and small sections of D-loop control region and 12S rRNA. Select annotations include genes indicated in green and rRNA indicated in red.

## Discussion

We demonstrated that with our NAS pipeline, we can successfully identify a species of an excreting host from a sample of their feces. We successfully sequenced sections of mtDNA, barcoding genes and/or complete or near-complete mitochondrial genomes of all of our samples. Eight out of nine identifications matched original PCR or morphology-based identification and we believe the ninth was originally misidentified by PCR due to a lack of supporting evidence from our study. Most methods for the molecular identification of mammal feces involve a PCR step that is time-consuming and prone to false negatives due to poor quality DNA. The NAS methods described in this paper can lead to an accurate and rapid (i.e., data generated within hours of collection) species identification completely free of PCR. Going from fecal sample to species identification quickly is invaluable to conservationists and biologists studying difficult to find or distinguish species by providing an accurate molecular-based identification in a timely manner. With faster methods for fecal identification, informed management plans or research can be carried out without waiting months for PCR results or visual sightings of species. This study serves as a proof of concept for this method and can be expanded to incorporate the generation of other sequencing data NAS targets.

In line with previous studies focused on mitochondrial based species identifications, as well as critiques of such methods, extra caution must be used when considering incomplete lineage sorting, mitochondrial capture, and historical or active hybrid zones (Larsen et al. 2010; Thompson et al. 2013; vonHoldt et al. 2016). In certain situations, using NAS for species identification may require extra care, for example when distinguishing particular wild species and their domestic counterparts (e.g., polecat and domestic ferret, wildcat and domestic cat, wolf, and domestic dog). Such examples are hampered by the sharing of mtDNA haplotypes as a result of incomplete lineage sorting or hybridization (Davison et al. 1999; Randi et al. 2001; Krofel et al. 2022). We found that using multiple bioinformatic tools in combination (i.e., minimap2, kraken2, Blast and BOLD search engines, phylogenetic tree generation) could sometimes help to parse out an identification in these situations (i.e., *C. lupus* versus *C. lupus familiaris*). Sample 8 was identified as *O. virginianus* by all bioinformatic methods, however in the phylogenetic tree generated by BOLD with COI sequences, the sample sequence grouped with sequences from both *O. virginianus* and *O. hemionus*. This result coincides with recent evidence supporting that these two species underwent a hybridization event approximately 1.32 mya. The resulting mitochondrial capture of the *O. virginianus* mitogenome by *O. hemionus* produced a new haplogroup, along with novel haplogroups produced from introgression during continuing hybridization events between the two species (Wright et al. 2022).

As with all molecular-based species ID analyses of bulk fecal samples, another limitation arises when attempting to differentiate between host and prey species, particularly among carnivores consuming mammalian prey. However, other characteristics of the feces sample being tested (e.g., morphological, ecological, temporal, etc.) could be used to elucidate host species.

For the Kraken2 analysis of sample 5, *L. rufus* was the match of the majority of reads, however, a small percentage of reads were mapping to *L. americanus* and *C. temmnickii. C. temmnickii* was ruled out due to only one read mapping to this species and geographical range of this species in relation to where our samples were collected. We could rule out a *L. americanus* identification for the depositing host because of the low number of reads mapping to this species compared to the number mapping to *L. rufus*. Fecal morphology and total size of the sample also supported the identification of *L. rufus* over *L. americanus*. However, *L. americanus* are an important prey animal of *L. rufus* (Moen et al. 2012). We note that extracting DNA from mucosal cells found at the exterior of fecal samples would likely enrich the depositing host species DNA and thus would be a useful approach in such situations.

While the methods described herein do not require PCR to produce species identifications, PCR amplification could be used to amplify mtDNA to increase sensitivity and produce more reads at higher coverage, although with the limitations described above. In light of our results, we recommend a minimum of 40 mitochondrial reads, having a minimum sequence length ranging from approximately 500 to 1,000 base pairs, for NAS-based species identification using scat samples. With these minimum specifications, it is more likely that a confident identification can be made based on the above methods. A Full-sized MinION flow cell (e.g., R10.4) clearly provides greater sequencing depth, thus yielding more mitochondrial host reads for species identification, than the smaller ONT Flongle. The current price for a full-size flow-cell is approximately $900 USD, though molecular barcoding and pooling individual samples can reduce cost per sample. The full-sized flow cells utilized for this project had all been used for other projects and then washed for reuse. Nevertheless, utilizing Flongles significantly reduces total cost (current price $90 USD) for NAS-based molecular barcoding. The flongles all had less than the maximum possible number of pores due to their age. The lowest pore count at the start of a flongle experiment leading to a successful identification was 24 out of a possible 126 pores. However, given our Flongle data, we recommend a minimum of at least 100 active sequencing pores at the start of an experiment to provide optimal results from individual samples. Even with these reduced pore counts, we still generated enough data to identify our samples, demonstrating another opportunity to reduce cost. We note that continual improvements to ONT sequencing technologies and associated bioinformatics will impact these estimates, lowering cost per sample and increasing sequencing throughput.

The resolving power of NAS-based species identification will increase concordantly with the increase in available mammalian mitochondrial genomes. At the time of this study, no mitochondrial sequences had been published to NCBI for *S. floridanus*. Because of this lack of comparative sequences, we made the species identification of sample 6 by mapping to closely related species and then using metadata including location of sample collection and species with typical ranges in that area. Sample and collection site metadata are an invaluable resource to validate molecularly generated identifications, especially when reference mitochondrial genomes of species of interest are not currently available.

We provide an accurate and quick procedure to identify mammalian species from a fecal sample. With our nanopore sequencing pipeline, from the time a fecal sample is acquired to bioinformatic analysis of DNA sequence data, a species identification can be achieved in less than 12 hours. The adaptive sampling method allows for fecal mtDNA to be enriched and targeted for sequencing which has previously required a PCR step. There is potential to use NAS to generate entire reference quality mitochondrial genomes while documenting microbiomes and diet components, such as consumed prey species and vegetation. Conducting in-the-field extraction and sequencing of DNA with a mobile lab, facilitated by the portability of the ONT MinION and miniaturization of required lab equipment, could provide even more flexibility and speed for molecular species identifications on site.

Future studies could apply this framework to detect barcoding genes and whole genomes of fecal pathogens. NAS of fecal samples for species identification can be easily applied to conservation, biodiversity studies, invasive species detection, illegal wildlife trade, marine mammal studies, and forensic studies that require fast and accurate species identification. Many North American Indian tribes are in the process of inventory and monitoring of biodiversity on tribal and ceded lands. The northeastern Ojibwe tribes authored in this study are all leading initiatives to measure biodiversity and map and document the spread of invasive species as climate adaptation plans are implemented (Moore et al. 2014; Stults et al. 2016). The tool described here will assist in those efforts. The ease of Nanopore library prep, requiring only basic pipetting skills and an understanding of the technology, and lack of PCR means that this molecular method is more accessible to many groups than PCR based methods.

## Acknowledgements

We thank Seth Stapleton and Minnesota Zoo Staff for providing access to zoo grounds and biological samples. We also thank the Leech Lake Band of Ojibwe, Red Lake Band of Chippewa Indians, and Grand Portage Band of Lake Superior Chippewa for generously contributing wild carnivore samples. LEF was supported by the K. V. Nagaraja Tuition Fellowship and by discretionary funds awarded to PAL during the course of the research. Startup funds awarded to PAL supported the overall research effort.

## Supplementary Data

**Supplementary Data SD1**. *—*Detailed bioinformatic pipeline including, software packages and versions, commands used for species identification, and explanation of usage.

**Supplementary Data SD2**. *—*Metadata from the neighbor joining phylogenetic tree generated by the BOLD database using the COI sequence from sample 8, identified as *Odocoileus virginianus*.

