## Supplementary Data 1 for "Rapid molecular species identification of mammalian scat samples using nanopore adaptive sampling": SD1_fecalID_bioinformaticpipeline.html

Mammal Species Identification from Feces using Nanopore Adaptive Sampling


### Mammal Species Identification from Feces using Nanopore Adaptive Sampling

###### Lexi Frank

#### 2023-03-28

Here is step-by-step instructions for the mammal scat species
identification pipeline.

#### Background

##### Experiment

Authors: Lexi E. Frank, Laramie L. Lindsey, Evan J. Kipp, Christopher
Faulk, Suzanne Stone, Tanya M. Roerick, Seth A. Moore, Tiffany M. Wolf,
Peter A. Larsen

Sample origins: Bernstein L.A., Shaffer C., Walz E., Moore S., Sparks
A., Stone S., Roerick T., Larsen P.A., Wolf T.M. 2021. Exploring risk
for echinococcosis spillover in northern Minnesota tribal communities.
EcoHealth. 18:169–181, the Minnesota Zoo, and opportunistic collection
within Minnesota.

Extraction method: Qiagen QIAmp PowerFecal Pro DNA kit

Sequencing instrument: Oxford Nanopore Minion, r9.4.1 flongle or
full-sized flowcell

Library prep: SQK-LSK109 or SQK-LSK110 kit.

Computational Environment: Linux desktop computer (Intel C600/X79
series i9-10920X 12 core; Linux 5.4.0-77-generic x86\_64; Ubuntu 18.04;
Nvidia Quadro RTX 4000 GPU with 8 GB video memory) or a Linux laptop
(16x 11th gen Intel Core i7; Ubuntu 18.04; Nvidia GeForce RTX 3080 Ti
GPU with 16 GB video memory)

#### Methods

##### Creating a Reference Database for Nanopore Adaptive Sampling

Download complete mitogenome RefSeq from NCBI
Organelle database.  
`wget https://ftp.ncbi.nlm.nih.gov/refseq/release/mitochondrion/mitochondrion.1.1.genomic.fna.gz gunzip *.fna.gz cat *.fna > mitogenome_refseq.fasta`

For faster computing and more accurate alignment of only mammalian
sequences, filter this file to only include mammals. Use NCBI Organelle
‘Browse
by organism’ function to pull mammal species names. Filter search by
SubGroup: Mammals and Type: mitochondrion. Under ‘Choose Columns’ select
Organism Name only; this will generate a table with all mammal species
names contained in the RefSeq; click download. Open the downloaded
organelles.csv file with a text editor. Remove the quotation marks with
search and replace function in a text editor and save as .txt file
(mammalnames.txt).

Some species may have multiple mitogenomes within the RefSeq and thus
appear twice in this .txt file; duplicate names can be removed using
sort.  
`sort -u mammalnames.txt > mammalnames_dedupe.txt`

Use seqkit to pull
NCBI header names out of mitogenome reference FASTA.  
`seqkit seq mitogenome_refseq.fasta -n > allmitoIDs.txt`

Use grep with both .txt files to pull header names corresponding to
mammals only.  
`grep -Fw -f mammalnames_dedupe.txt allmitoIDs.txt > allmammalIDs.txt`

Use filter by name script from BBMap
package to pull mitogenome sequences for mammals out of the complete
RefSeq FASTA.  
`/home/lexi/anaconda3/envs/bbmap/bin/filterbymap.sh in=mitogenome_refseq.fasta out=mammal_mitogenomes.fasta names=allmammalIDs.txt include=t`

Check the output file to ensure only mammal sequences are included.
Use this file as input into MinKNOW software for adaptive sampling and
real-time alignment.

##### Read Generation

Fast5 files were generated from the Oxford Nanopore MinION sequencer
and basecalled with Guppy v5.0.11. For post-hoc basecalling, we used the
super accuracy model with the following post-hoc parameters.  
`guppy_basecaller --config dna_r9.4.1_450bps_sup.cfg --device cuda:0 -q 0 --disable_qscore_filtering --nested_output_folder --recursive --input_path nosample/ --save_path sup_reads/`

#### Analysis

Combine fastqs generated from a single sample during basecalling into
1 fastq so they’ll be easy to work with.  
`cat fastq/*.fastq > allnano020.fastq`

##### Filtering for Quality and Length

Use nanofilt to
filter the reads for quality and length. We are targeting the
mitochondrial genome, which is around 16kb in length in mammals. So we
filter for reads that are between 300 and 17,000kb.  
`NanoFilt -q 7 -l 300 --maxlength 17000 allnano020.fastq > nano020_q7_300_17k.fastq`

##### Map Reads to reference database

We mapped our sample’s reads to our reference database using minimap2. Here our reference
database is the NCBI mammal mitochondrial RefSeq file created above (See
Creating a Reference Database for Nanopore Adaptive Sampling).  
`minimap2 -ax map-ont mammal_mitogenomes.fasta nano020_q7_300_17k.fastq > nano020_mitoreads.sam`

Use samtools to manipulate files. Change the .sam file to a .bam
file.  
`samtools view -bS nano020_mitoreads.sam > nano020_mitoreads.bam`

Reorganize the order of the .bam files to match the order of the
genome/gene.  
`samtools sort nano020_mitoreads.bam -o nano020_mitoreads_sorted.bam`

Pull out reads that mapped to the reference.  
`samtools view -h -F 4 -b nano020_mitoreads_sorted.bam > nano020_mapped.bam`

Change .bam file to .fastq file format.  
`samtools bam2fq nano020_mapped.bam > nano020_mapped.fastq`

##### Assemble the Mitochondrial Genome

Here we assemble the entire mitochondrial genome using the Flye assembler. If the
entire mitochondrial genome is not able to be assembled, move on to the
Kraken2/Pavian section.

Use reads that mapped to the reference database file in Flye to
assemble the mitogenome.  
`flye --nano-corr nano020_mapped.fastq --genome-size 16k --out-dir Assembly_Nano020`

Check the assembly by opening the assembly\_info.txt to see if the
mitogenome was assembled. Look for a single contig around 16kb in length
and usually circular (but not necessarily). Check the coverage (higher
coverage is better, 30X coverage considered to be high quality). Once
you determine which contig is likely the mitogenome, open the
assembly.fasta file. Copy and paste the sequence into NCBI
Blast to search for matches to the contig.  
`cd Assembly_Nano020`

`nano assembly_info.txt`

`cat assembly.fasta`

##### Annotate the Mitochondrial Genome

Go to Mitos2. Fill out
form to submit a mitogenome using Ref89 Metazoa and Vertebrate Code. You
can wait for the analysis to finish or they will email you when it is
done. Download annotation fasta after that analysis finishes. Open using
a text editor. Find the COI gene. Copy the sequence. Go to the BOLD
COI database, paste sequence and run search to find best matches. The
same can be done with NCBI Blast search and the cytb gene.

If no mitogenome was assembled, that is okay. The reads that mapped
to the mitogenome reference file (called nano020\_mapped.fasta, above)
can be used to generate an ID.

#### Species ID

At this point, a species ID can be achieved in several ways depending
on the quality and coverage of the sequencing data and whether or not a
full mitogenome was assembled.

##### Kraken2/Pavian

Build a Kraken2
database with the NCBI mitochondrial refseq file then use Kraken2 to map
the reads to this database using a k-mer-based approach to provide
taxonomic classifications of sequences.  
`kraken2-build --download-taxonomy --db Mammal_scat`

`kraken2-build --download-library Univec --db Mammal_scat`

`kraken2-build --add-to-library mitochondrion.1.1.genomic.fna --db Mammal_scat`

`Kraken2-build --build --db Mammal_scat --threads 8`

Run mapped sample reads against the kraken2 database.  
`kraken2 --db Mammal_scat --threads 8 nano020_mapped.fastq --report nano020_mapped.txt`

Launch Pavian in
R studio to visual the Kraken2 data.  
`pavian::runApp(port=500)`

Load the report file, nano020\_mapped.txt, into Pavian to view figure.
Ensure that the proper filters are in place in the taxon filter at the
top. Once filtered, the mammal species with the most reads mapping to it
is likely the correct ID. However, if the species mitogenome has not
been published yet or was not in the NCBI refseq database then several
closely related species may have reads matching to them. This method
alone cannot necessarily distinguish between domestic species and their
wild counterparts (i.e. wolf and dog) or species with mitochondrial
capture/hybridization. Move on to phylogenetic tree generation for more
information and utilize additional metadata (location of sample
collection, geographic species ranges, size and morphology of feces) to
differentiate in these situations.

##### Phylogenetic Tree Generation

###### Geneious Software

Use the reference genome from NCBI of the best-guess or putative
species ID of the sample and align it to the sample’s mapped reads fastq
file in Geneious Prime software
(v2022.2.1). This will allow you to find where genes are located within
the mitogenome or determine which reads are covering the barcoding genes
(cytb or COI). OR Annotate in Geneious with the MITOS2 output Download
the .GFF output file from Mitos2. Convert the file to a Genbank file
using Emboss
seqret.  
`seqret -sequence contig_56_mitogenome.fasta -feature -fformat gff -fopenfile contig56mitogenome.gff -osformat genbank -auto`

Bring the genbank file into Geneious.

A phylogenetic tree can be created in Geneious using RAxML or
in-command line RAxML. In Geneious, gather sequences of entire
mitogenomes, cytb, or COI from several different species closely related
to the suspected ID of the sample. Multiple align these sequences and
then use RAxML to generate the phylogenetic tree with GAMMA model with
1,000 bootstrap iterations.

###### RaXML Phylogenetic Tree without Geneious software

Create a phylogeny based on either cytb gene or COI gene. Use the
gene sequence from Mitos2 for cytb or COI. Make a file with sample
sequence and sequences of closely related species from NCBI.  
`nano mammal_gene.fasta`

Use mafft to align sequences.  
`mafft mammal_gene.fasta > COI_Aln.fasta`

Use Fasta2Phylip
perl script to change format of alignment to phylip.The phylogenetic
program we use will want a phylip format.  
`perl Fasta2Phylip.pl COI_pero_Aln.fasta COI_pero_Aln.phy`

RAXML is the phylogenetic analysis program we want to use to create a
tree.  
`raxml -f a -m GTRGAMMAI -p 12345 -x 98765 -s COI_Aln.phy -n COI_1000 -# 1000 -o Sample`

To view the tree, use the bipartitions file created from raxml. Use
FigTree. Add
Node labels and use drop down menu to select ‘label’. Adjust the size of
tree, branch labels, etc. Export graphic as an enhanced meta file (.emf)
which can be edited in powerpoint or as an svg which can be edited in Inkscape. Insert the file, ungroup the
image and edit labels, branches, etc.

###### BOLD database

Copy and paste the consensus sequence of the COI barcoding gene from
Geneious software into the BOLD
search engine. This generates an id based on similarity to the BOLD
database of COI sequences and a phylogenetic neighbor-joining tree.

###### NCBI blast

Copy and paste the consensus sequence of the COI/cytb/mitogenome
sequence from Geneious software into the Blast
search engine. This generates an id based on similarity to the Blast
database of sequences and a phylogenetic neighbor-joining tree.

#### Nanopore Experiment Metadata

To view information on the adaptive sampling run, go to the adaptive
sampling .html
/home/plarsen/shared/Mammal\_Scat/Mammal\_scat\_nano011\_8Aug2022/no\_sample/20220808\_1043\_MN27245\_AJL698\_24bdeb57/
report\_AJL698\_20220808\_1044\_24bdeb57.html

To generate files on the stats from the run, we ran nanoplot and then
looked at the .html file that is generated.  
`NanoPlot -t 2 --fastq allnano020.fastq`

To generate .html file on other stats from the run, we used fastqc.  
`fastqc allnano020.fastq --out fastqc_out`
